# Angiocrine IGFBP3 Spatially Coordinates IGF Signaling During Neonatal Cardiac Regeneration

**DOI:** 10.1101/2021.09.16.460522

**Authors:** Shah R. Ali, Waleed Elhelaly, Ngoc Uyen Nhi Nguyen, Shujuan Li, Ivan Menendez-Montes, Zhaoning Wang, Miao Cui, Abdallah Elnwasany, Feng Xiao, Suwannee Thet, Nicholas T. Lam, Alisson Cardoso, Ana Helena Pereira, Mohammad Goodarzi, Michael T. Kinter, Andrew Lemoff, Luke I. Szweda, John Shelton, Wataru Kimura, Hesham A. Sadek

**Author notes:** Authors of correspondence, Corresponding Authors: Shah R. Ali MD, Hesham A. Sadek MD, PhD.

## Abstract

To identify non-cell-autonomous effectors of cardiomyocyte mitosis, we analyzed a transcriptomic screen of regenerating and non-regenerating neonatal hearts for differentially-expressed secreted proteins – which we hypothesized could include candidate mitogens. We identified and validated IGFBP3, which has a Janus-like stabilizing and sequestering effect on IGF growth factors, as a neonatal injury-associated secreted protein. IGFBP3 is expressed by and secreted from vascular cells in the neonatal heart after cardiac injury, notably in the infarct border zone. We found that global deletion of IGFBP3 blunted neonatal regeneration, while gain-of-function experiments using recombinant IGFBP3 and a transgenic mouse model uncovered a pro-mitotic effect of IGFBP3 on cardiomyocytes in vitro and in the adult heart. We show that site-specific expression of an IGFBP3 protease (PAPP-A2) and its inhibitor (STC2) coordinate the spatial release of IGF2 in the infarct zone to regio-selectively activate the INSR-ERK-AKT cell growth pathways in cardiomyocytes. Collectively, our work highlights the spatiotemporal orchestration of endothelial-cardiomyocyte interactions that are required for neonatal cardiac regeneration.

## Introduction

It is well-recognized that the adult mammalian heart can only meagerly regenerate after a myocardial infarction (MI) injury, which results in the replacement of functioning myocardium with a fibrotic scar.^1^ The prevalence of MI and its sequelae, e.g., heart failure, are a major contributor to the high burden of morbidity and mortality posed by cardiovascular disease across the globe.^2^ The mechanistic basis for the heart’s paltry regenerative response upon injury resides in the cardiomyocyte: it is unable to meaningfully divide in the adult mammalian heart.^3–5^ As such, significant efforts are underway to decipher why the cardiomyocyte cloisters its cell cycle; identifying novel strategies to inuce cardiomyocyte mitosis – to regenerate the adult heart – represents a new paradigm for the treatment of heart diseases in humans.^6^

Unlike the adult, the neonatal heart in many mammalian species can regenerate after various insults, including MI.^7–11^ Neonatal cardiac regeneration provides evidence that postnatal cardiomyocytes can divide during a discrete temporal window. We hypothesized that study of this phenomenon could yield molecular insights about the extrinsic factors and signaling pathways that underlie myocyte proliferation. Although the transcriptional programs that occur during neonatal regeneration *in toto* and in individual cells have been previously annotated, how individual proteins from different lineages interact to orchestrate the complex regenerative program seeks to be defined.^12–15^

Recent studies have highlighted a wide range of mechanisms that undergird neonatal heart regeneration.^16–27^ Of interest to the current report, the critical role of the IGF signaling pathway in murine neonatal cardiac regeneration was recently established through loss-of-function experiments of its key ligands and receptors; previous studies meticulously laid the groundwork for these findings by demonstrating that the same pathway is ensconced in murine embryonic cardiac development and zebrafish adult cardiac regeneration.^28–32^ While they revealed key insights, these studies also generated new questions about how these signals are elaborated and coordinated after injury to impact the regenerative response. For example, while both the IGF2 ligand and its cognate receptor are expressed after neonatal and adult MI, the receptor is activated only after neonatal MI. Such results corroborate prior attempts to renew the adult heart with recombinant or adenoviral delivery of IGF ligands that showed modest beneficial outcomes.^33–35^ These experiments directly imply the existence of a sub rosa, upstream regulatory schema that might reconcile these discordant findings between young and old hearts.

In this work, we characterize a spatiotemporally coordinated paracrine signaling axis involving vascular cells and cardiomyocytes that regulates IGF signaling to elicit cardiomyocyte proliferation, which is the *sine qua non* of neonatal cardiac regeneration. By elucidating a pro-regenerative module constituted by the intercellular crosstalk among distinct cell lineages, these results highlight the strict regulation of IGF pathway components across space and time during neonatal regeneration and offer a potential therapeutic path to modify the regenerative potential of the adult heart.

## Results

### IGFBP3 is expressed in vascular cells in the heart after neonatal MI

We sought to identify mitogenic secretory proteins that help elicit myocyte proliferation in an unbiased manner. We performed a transcriptomic screen on ventricular tissue from regenerating (P1) and non-regenerating (P14) hearts 3 days after MI (by ligation of the left anterior descending artery (LAD)) or sham operation using microarrays (n=3 for each group). We analyzed this data by focusing on the expression levels of genes annotated as conventionally secreted (Fig. 1A). Imposing these bioinformatic restrictions revealed 149 genes encoding putative secreted proteins (Fig. 1B, Supplementary Table 1). We hypothesized that candidate secretory genes would likely be upregulated after neonatal MI compared to sham or non-regenerative MI. Several genes followed this expression pattern, e.g., Igfbp3, Angpl7, which we further studied (Fig. 1C). To scrutinize the expression patterns of these genes at the tissue level, we performed in-situ hybridization (ISH) on post-operative day 3 (POD3) tissue after regenerative (P1 MI) and non-regenerative (P7 MI). Of these, only Igfbp3 ISH recapitulated its microarray pattern (Fig. 1G-J, Fig. S1): namely, Igfbp3 transcript was upregulated after neonatal MI and sparsely detected after P7 MI. More, the expression pattern of Igfbp3 was prominently injury-associated: after neonatal MI, Igfbp3 transcript was localized to the border zone surrounding the infarct (denoted by the dearth of Troponin T ISH staining) (Fig. 1G-H). This suggested that Igfbp3 is expressed at the right time and place to be a putative pro-mitotic secreted gene with a role in neonatal regeneration. Therefore, we further characterized the role of Igfbp3 following MI.

**Figure 1:**
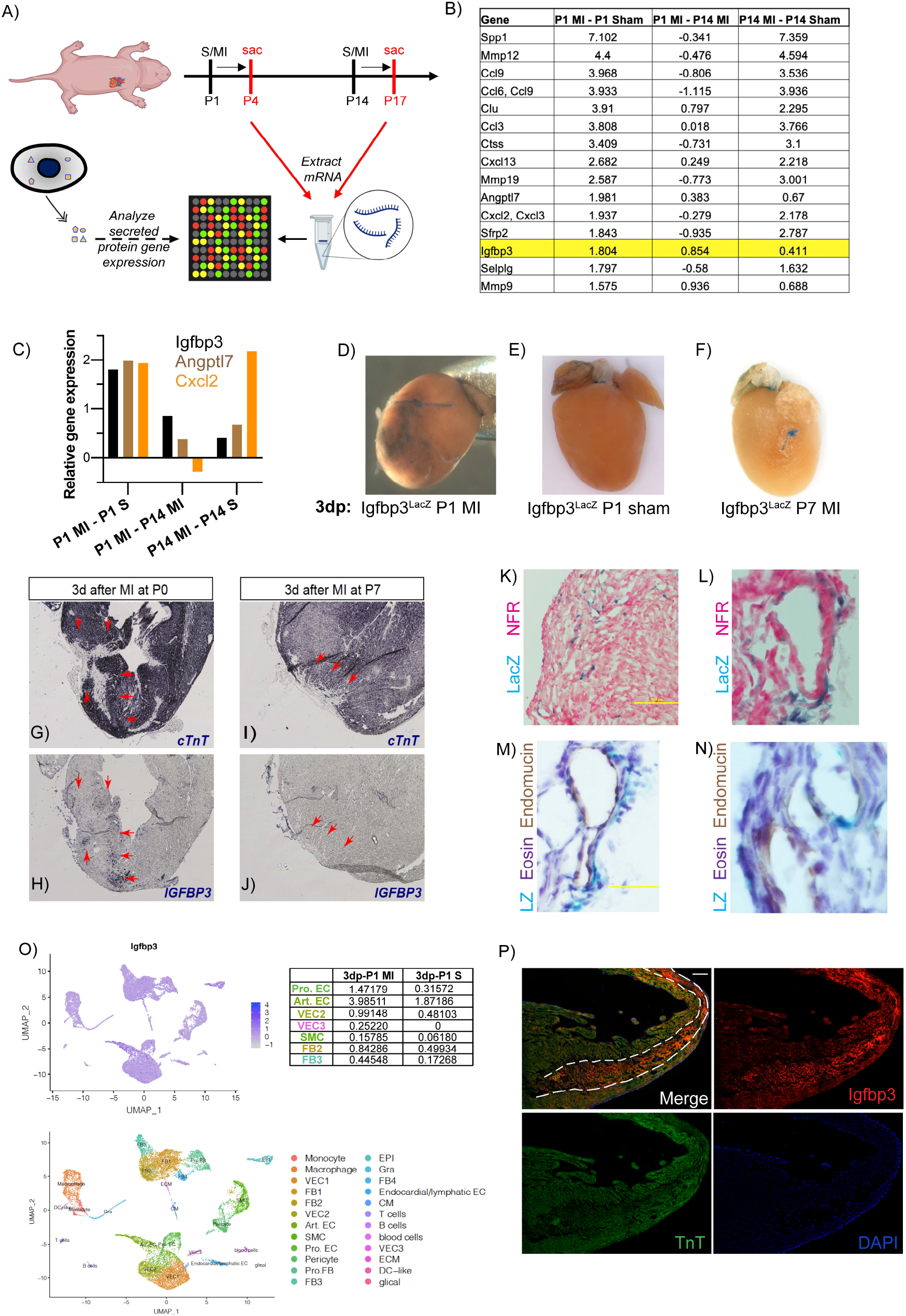
Igfbp3 is expressed in the heart after neonatal MI. A) Schematic depicting experimental approach. B) Curated list of candidate secreted genes based on bioinformatic analysis after neonatal and adult MI or sham. C) Relative gene expression of discrete secreted genes after MI or sham. D-F) LacZ staining of Igfbp3^LacZ^ reporter mice 3 days post (3dp) specified operation and timepoint. G-J) In situ hybridization of cardiac tissue 3 days after MI at P0 (G-H) or P7 (I-J) for cardiac Troponin T (cTnT) (G,I) or Igfbp3 (H,J). Red arrows outline infarct border. K-L) LacZ staining (and nuclear fast red (NFR) counterstain) of cardiac sections 3 days after P1 MI. M-N) LacZ staining, eosin counterstaining, and endomucin immunohistochemistry on cardiac sections 3 days after P1 MI. O) Igfbp3 scRNA-seq of cardiac cells after P1 MI (data derived from Wang et al.) Table includes scRNA-seq gene expression values of Igfbp3 in TPM (total counts per million) in various cell lineages (color-coded) from cardiac tissue 3 days after P1 MI or sham (Art.EC, arterial endothelial cell; Pro.EC, proliferating endothelial cells; Pro.FB, proliferating FB; SMC, smooth muscle cell; VEC, vascular endothelial cell.) P) Immunofluorescent staining of cardiac tissue 3 days after P1 MI. Dashed white line demarcates infarct boundary. Scale bar 100 μm.

We used an Igfbp3^LacZ^ reporter mouse line, in which LacZ is expressed in an Igfbp3-dependent manner (in place of the native Igfbp3 allele, leading to loss of function) (Fig. 2A).^36^ We performed whole-mount LacZ staining on POD3 after P1MI (or sham) and after P7 MI, which showed patchy LacZ staining diffusely below the surgical ligature after P1 MI (Fig. 1D). In contrast, only sparse atrial and great vessel staining was observed after P1 sham or P7 MI (Fig. 1E-F). These data collectively validate the initial screen findings to indicate that: 1) Igfbp3 mRNA is expressed in the heart uniquely after neonatal MI (compared to aging or non-regenerative injury), and 2) its expression is prominent in injury-associated tissue.

**Figure 2:**
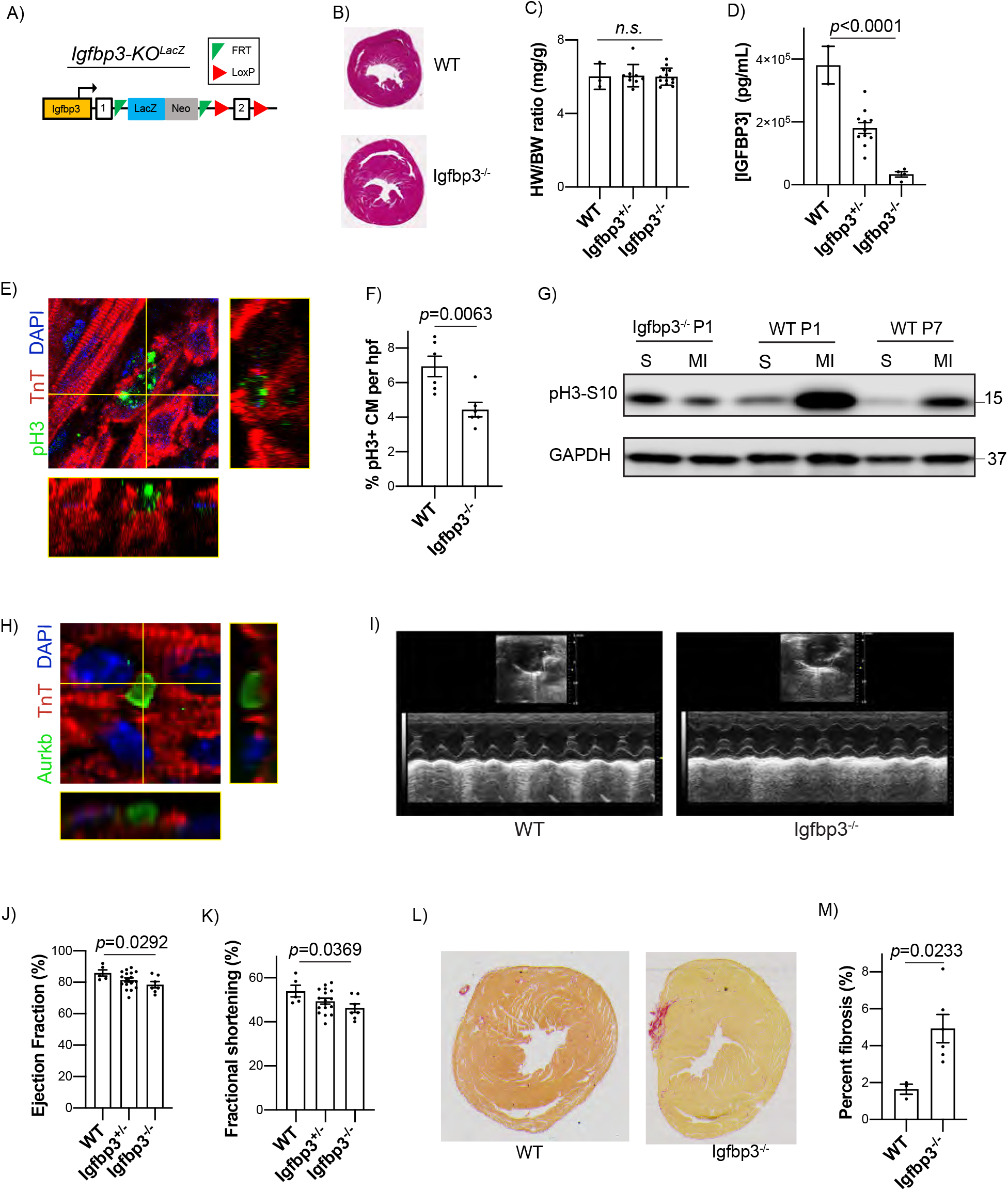
Igfbp3 deletion impairs neonatal regeneration. A) Genetic construct of Igfbp3-KO^LacZ^ model. B) H&E-stained short axis sections from one-month-old hearts. C) HW/BW ratio (mg/g) from one-month-old hearts. D) IGFBP3 protein levels (pg/mL) in serum determine with ELISA. E) Confocal microscopic image of an immunofluorescently-stained pH3+ cardiomyocyte (yellow lines correspond to z stack axes). F) Quantification of pH3+ CM per high-powered field (hpf) (n=6 animals per group). G) Western blot of pH3-S10 and GAPDH (loading control) from cardiac ventricular tissue 3 days after described operation and genotype (representative sham and MI displayed). H) Confocal microscopic image of inter-myocyte Aurora B kinase immunofluorescent staining (yellow lines correspond to z stack axes). I) Representative M-mode echocardiographic mid-systolic images at the level of the papillary muscle from hearts 4 weeks after P1 MI. J,K) Systolic function 4 weeks after P1 MI. L) Representative short-axis Picosirius Red-stained cardiac sections 4 weeks after P1 MI below level of ligature. M) Fibrosis quantification 4 weeks after P1 MI using Picosirius Red staining.

To delineate which cardiac cells express Igfbp3 during neonatal regeneration, we examined LacZ-stained tissue sections. These revealed that LacZ was often intercalated with blood vessels. We therefore performed immunohistochemistry on LacZ sections for endomucin, which is an endothelial-specific transmembrane protein (Fig. 1M-N).^37^ LacZ and endomucin were intimately co-localized, reinforcing a vascular origin for Igfbp3 after neonatal MI. We also analyzed a reported single cell RNA sequencing (scRNA-seq) repository from neonatal MI tissue, which showed that endothelial cells and fibroblasts are the two highest Igfbp3-expressing lineages in the regenerating neonatal heart post MI, providing another line of evidence for blood vessels expressing and secreting Igfbp3.^14^

The data so far evaluated the expression of Igfbp3 at the mRNA level, but Igfbp3 is known to be a secreted protein. Therefore, we sought to characterize whether IGFBP3 protein is found in the same location as its mRNA. We performed immunofluorescent staining on P4 cardiac sections after P1 MI, which strikingly showed that IGFBP3 is present within the infarct zone (Fig. 1P, Fig. S3), in stark contrast to the expression of Igfbp3 mRNA in the injury border zone (Fig. 1H). These findings suggest a model in which vascular cells, especially those proximal to the infarct region, express Igfbp3 – possibly in response to hypoxia, a known activator of Igfbp3 – which subsequently travels to the infarct zone.^38, 39^

Igfbp3 was originally identified as a plasma protein that stabilizes IGF ligands in circulation, although our findings stem from its cardiac-specific expression in response to a cardiac injury.^40^ IGF-as well as non-IGF-related functions of IGFBP3 have been well-characterized, which are pleiotropic, context-specific, and range from antiproliferative to pro-apoptotic to pro-proliferative.^41–44^ Since the rich body of work on Igfbp3 did not obviate its consideration as a pro-regenerative secreted factor, and because its expression pattern in the heart after injury showed a distinct regeneration-associated signature, we proceeded to fully interrogate the role of Igfbp3 during neonatal regeneration.

### IGFBP3 Deletion Impairs Neonatal Regeneration

We generated global Igfbp3 knockout mice (Igfbp3^-/-^), which are born to term at Mendelian ratios and exhibit grossly normal cardiac morphology (Fig. 2A-B).^36^ We chose to use a global, rather than a conditional deletion mouse model – which is achievable with the targeting strategy – given the expression of Igfbp3 in at least two cell types following MI: thus, targeted deletion in a single lineage is unlikely to eliminate Igfbp3 protein. The knockout mice had nearly undetectable levels of IGFBP3 in the serum (Fig. 2C). In addition, Igfbp3^-/-^ mice have a comparable heart weight-to-body weight ratio as WT littermate controls and show normal cardiac function (Fig. 2B, Fig. S2A). This suggests that Igfbp3 is dispensable for cardiac development and maturation, which aligns with the bland expression pattern of Igfbp3 without neonatal injury (Fig. 1E); in contrast, a related family member, Igfbp4, was found to be necessary for cardiogenesis in *Xenopus*.^45^ Since Igfbp3 was discovered in our screen as a putative secreted mitogen with robust expression after P1 MI, we speculated that any consequences of the knockout may only be revealed after neonatal injury.

Therefore, we performed P1 MI on Igfbp3^-/-^ pups and littermate controls (WT) to determine if deletion of this secreted protein could perturb cardiac regeneration. We first quantified the degree of cardiomyocyte proliferation at POD7, a time point known for robust cardiomyocyte renewal during the regenerative window after neonatal MI. Fewer myocytes underwent cell division using phosphorylated Histone H3 (pH3) as a marker of mitosis (7 pH3+ CMs vs. 4.5 pH3+ CMs per section, p=0.0063) (Fig. 2E-F). We also evaluated if Igfbp3 loss of function affects cardiomyocyte cytokinesis using Aurora kinase B immunofluorescent staining: we found that compared to WT, fewer cardiomyocytes exhibited inter-myocyte Aurora B kinase expression in the knockout hearts (Fig. 2H). We next studied the functional effect of Igfbp3 loss after P1 MI using echocardiography at 1 month after injury. Compared to WT mice, which achieved normal cardiac function, Igfbp3^-/-^ mice show incomplete recovery of cardiac function (ejection fraction (EF) 86% vs. 78%, p=0.0292) and fractional shortening (FS) 54% vs 46%, p=0.0369) (Fig. 2I-K). We subsequently explanted the hearts 1-month post-injury and quantified the size of the fibrotic scar (using Picosirius Red staining), which is present after adult MI but absent after neonatal MI. We found that WT hearts show nearly no Picosirius red staining (1.6% fibrosis), as previously described, but Igfbp3^-/-^ hearts had 4.9% fibrosis (p=0.0233) (Fig. 2L-M). These data collectively indicate that although the deficit of Igfbp3 has no discernible effect on cardiac development and physiology, it reduces cardiomyocyte proliferation and impairs cardiac regeneration after P1 MI.

### Recombinant IGFBP3 Promotes Cardiomyocyte Proliferation *In Vitro* and *In Vivo*

Given that Igfbp3 is necessary for cardiomyocyte renewal during neonatal regeneration, we sought to determine whether it is sufficient to elicit cardiomyocyte replication. We first utilized an *in vitro* model of neonatal rat ventricular myocytes (NRVM) in culture. We initially attempted to express IGFBP3 in a bacterial system, which was unsuccessful due to protein aggregation (data not shown). Given the complex post-translational modifications of IGFBP3, which may not be feasible in bacteria, we generated a stable 293T HEK cell line lentivirally infected with a C-terminus FLAG epitope-tagged mouse IGFBP3 (Fig. 3A). We confirmed that this transgenic cell line ectopically expresses Igfbp3 mRNA and secretes IGFBP3-Flag protein into the conditioned medium (Fig. 3B-D). We were able to isolate the IGFBP3-Flag using magnetic Flag antibody beads for further characterization. The band for the Flag immunoblot on the purified, denatured protein was consistently ~10-15 kDa higher than the predicted molecular weight of mouse IGFBP3 (32kDa) (Fig. 3E). We suspected that this may be due to post-translational glycosylation, for which IGFBP3 has 3 predicted asparagine residues.^40, 46^

**Figure 3:**
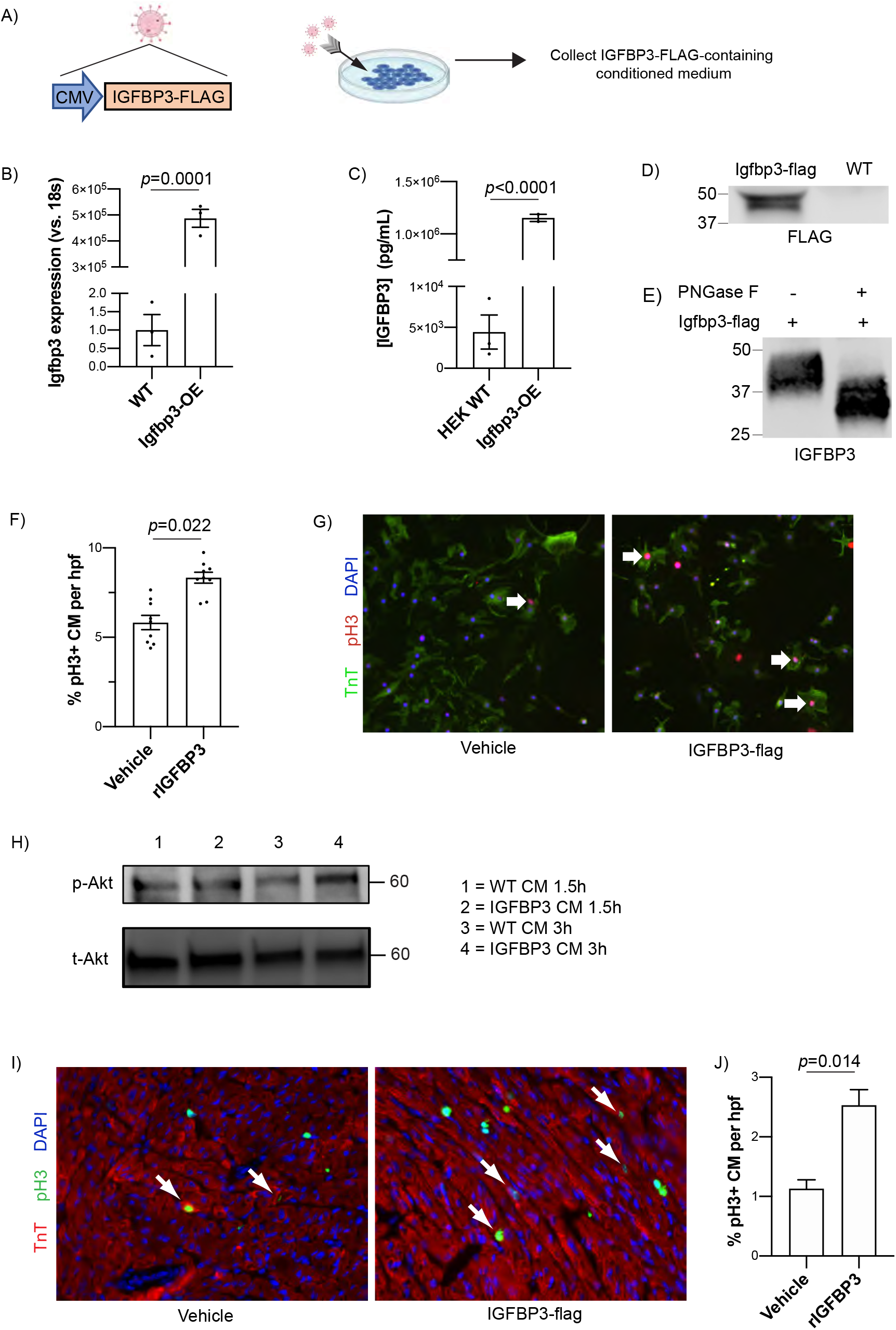
Recombinant IGFBP3 promotes cardiomyocyte proliferation in vitro and in vivo. A) Genetic lentiviral construct for recombinant Igfbp3-flag (rIGFBP3) expression in 293T cells and schematic outlining experimental approach. B) IGFBP3 protein levels in the conditioned medium of 293T cells (pg/mL) determined with ELISA. D) Western blot for Flag from cellular lysates of 293T cell lines. E) Western blot for IGFBP3 on purified rIGFBP3-Flag protein with or without treatment with PNGase F. F) Quantification of pH3+ NRVM per high-powered field (hpf) with or without incubation with rIGFBP3. G) Epifluorescent microscopic images of NRVM co-incubated with vehicle or with rIGFBP3. H) Western blot of phosphorylated Akt (p-Akt) and total Akt (t-Akt) from cell lysates of NRVM incubated with wild-type conditioned medium (CM) (lanes 1,3) or IGFBP3-overexpressed CM (lanes 2,4) at 1.5 hours (h) or 3h after incubation. I) pH3 immunofluorescent staining of cardiac sections 2 days after intracardiac injection of vehicle or rIGFBP3. Arrows indicate pH3+ CM. J) Quantification of pH3+ CMs in cardiac sections from intracardiac-injected rIGFBP3 or control hearts.

We treated the purified IGFBP3-flag protein with PNGase F, a deglycosylase, which shifted the size to the predicted weight (Fig. 3E).^47^ Additionally, treatment with the N-glycosylation inhibitior tunicamycin appropriately decreased the molecular weight of the secreted IGFBP3-Flag protein in the conditioned medium (Fig. S4A).^48^

We then tested whether secreted Igfbp3 can induce cardiomyocyte proliferation in vitro. Culturing NRVMs in conditioned medium from wildtype HEK cells showed a basal proliferation rate of 5.8% cardiomyocytes (that express pH3 by immunofluorescence). Incubation of NRVM in medium supplemented with rIGFBP3 increased the fraction of mitotic NRVM to 8.3% pH3+ NRVM (p=0.0022) (Fig. 3F-G). We coimmunoprecipitated IGFBP3-Flag incubated with NRVM, which detected IGF2; IGF1 was not detected (Supplementary Table 2). This finding suggests that IGFBP3 affects NRVMs through the canonical IGF pathway^49–51^. Accordingly, phosphorylated AKT (pAKT) levels were higher in rIGFBP3-treated NRVM (Fig. 3H), which is a known downstream target of the IGF signaling pathway^52, 53^

We subsequently injected recombinant IGFBP3 into the mouse heart at P7, a postnatal time point when there is limited cardiomyocyte division during physiological homeostasis and limited myocardial regenerative response following injury. Compared to saline control-treated hearts, rIGFBP3-treated hearts had a greater fraction of mitotic cardiomyocytes (1.3 vs. 2.6 pH3+ cardiomyocytes per high-powered field, p=0.035) two days after injection (Fig. 3I-J). The outcome of these experiments thus suggested that rIGFBP3 may be able to promote cardiomyocyte proliferation both in cell culture and in the postnatal mouse heart.

### Generation of a Cre-inducible Igfbp3 ectopic expression mouse model

We speculated that different cell lineages may offer disparate post-translational modifications of IGFBP3, and that the rIGFBP3 expressed in a human cell line (293T) may not phenocopy the effects of tissue-expressed IGFBP3 in the mouse heart. Therefore, we generated a novel conditional, ectopic Igfbp3 transgenic mouse model, in which Cre induction would lead to expression of mouse Igfbp3 and GFP in a monocistronic manner (Fig. 4A). We knocked this cassette, driven by a CAG promoter, into the Rosa26 locus using CRISPR-Cas9 genome editing using blastocyst transgenesis.^54^ We obtained 3 transgenic mouse lines, of which 2 were knocked into the Rosa26 locus, as confirmed by PCR (Fig. S4B). We cultured primary ear-tip fibroblasts from the knock-in lines and transduced them with Cre lentivirus to delete the floxed stop codon and activate Igfbp3 expression (Fig. 4B). We validated that the knock-in mouse-derived fibroblasts had more robust secretion of IGFBP3 into the conditioned medium compared to control fibroblasts using ELISA (Fig. 4C). Interestingly, even wild-type cells demonstrated IGFBP3 in the conditioned medium, which we postulated was derived from the fetal bovine serum-supplemented basal medium. Having validated the transgenic model in culture, we proceeded to cross it to a Cre-bearing model.

**Figure 4:**
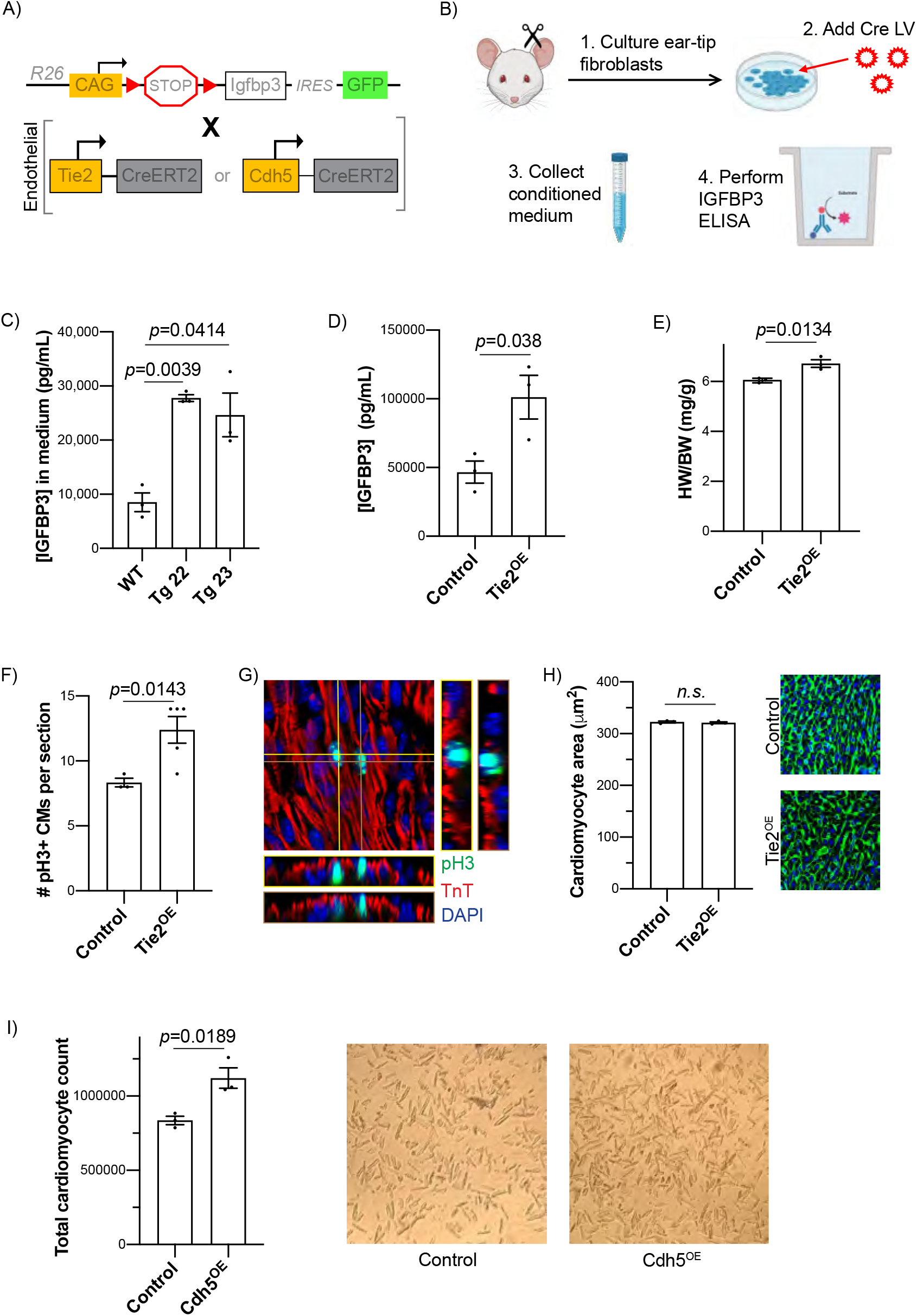
Transgenic IGFBP3 expression stimulates postnatal cardiomyocyte mitosis in vivo. A) Genetic model of knock-in Igfbp3-overexpression model and crosses with endothelial-specific Cre drivers. B) Schematic depicting approach to validate Igfbp3-overexpression mouse model using an in vitro CM-based ELISA assay. C) IGFBP3 protein levels (pg/mL) from conditioned medium of primary mouse ear-tip fibroblasts derived from two knock-in transgenic lines. D) IGFBP3 protein level (pg/mL) from serum determined with ELISA (n=3 per group). E) HW/BW ratios (mg/g) of P7 mice (n=3 per group). F) Quantification of pH3+ cardiomyocytes per section from P7 mice (n=3 for control; n=5 for Tie2^OE^). G) Representative confocal microscope image of cardiac section immunostained against pH3 and TnT from Tie2^OE^ heart (yellow and brown lines correspond to respective z stack axes). H) Cell size quantification using WGA staining from P7 hearts and representative confocal images. I) Total number of cardiomyocytes per heart after digestion (n=3 per genotype), with representative images from bright-field microscope.

### Endothelial-Specific Ectopic IGFBP3 Expression Can Regenerate the Postnatal Mouse Heart

While several lineages expressed IGFBP3 after P1 MI, we focused on endothelial cells given that angiocrine factors are known to mediate regeneration in various organs.^55–57^ Therefore, we used a Tie2-CreERT2 transgenic mouse line, which labels predominantly endothelial cells (with some contribution to hematopoietic lineages), to ectopically express Igfbp3 (Tie2^OE^) (Fig. S4C).^58^ We first activated Cre in the first week of life, when there is a small degree of basal cardiomyocyte replication, and studied the effects of Igfbp3 overexpression at P8. The transgenic mice had a higher heart weight-to-body weight ratio compared to littermate control mice (6.7 vs. 6.0 mg/g, p=0.0134) (Fig. 4E), suggesting cardiomyocyte hyperplasia or hypertrophy. We analyzed these mice for markers of cardiomyocyte mitosis, and we found an increase in pH3+ CMs in the Igfbp3^OE^ mice compared to the controls (12.4 vs. 8.3 pH3+ CMs per section) (Fig. 4F-G); we did not observe a change in cardiomyocyte cell size (Fig. 4H). This indicates that, during the neonatal window of cardiomyocyte proliferation, ectopic endothelial-derived IGFBP3 can increase the number of cycling myocytes. We also utilized an additional endothelial Cre line, Cdh5-CreERT2, and we administered tamoxifen during week 5-6 of adult life.^59^ At week 9, we digested the hearts and isolated cardiomyocytes: control hearts had, on average, 0.84 million cardiomyocytes compared to 1.1 million cardiomyocytes for the Cdh^OE^ hearts (p=0.0189), further reinforcing the mitotic effect of ectopic IGFBP3 on postnatal cardiomyocytes (Fig. 4I).

### Spatiotemporal Expression of IGFBP3 Regulators Occurs After Neonatal MI

We reasoned that the mechanism of action of IGFBP3 during neonatal regeneration relies upon canonical IGF signaling based on the NRVM coIP (Supplementary Table 2). Therefore, we performed immunoblots for IGF pathway components by immunoblot at POD3 after P1 MI (or sham) in WT and Igfbp3^-/-^ hearts and in WT P7 MI (or sham) hearts (Fig. 5A, Fig. S5A-G). We detected robust phosphorylation of IGF1-R/INS-R in WT tissue after P1 MI compared to P1 sham, a change that was not observed after P7 MI and P7 sham. Intriguingly, receptor activation is markedly attenuated in Igfbp3^-/-^ tissue after P1 MI (Fig. 5A, Fig. S5G). Concordantly, downstream targets of the IGF pathway – such as Akt and Erk1/2 – also exhibit blunted activation after WT P7 MI and Igfbp3^-/-^ P1 MI, in concert with abrogated activation of the upstream receptors (Fig. 5, Fig. S5A-F). These data demonstrate the salience of IGF signaling for neonatal heart regeneration and the central role of IGFBP3 in effecting the IGF response.

**Figure 5:**
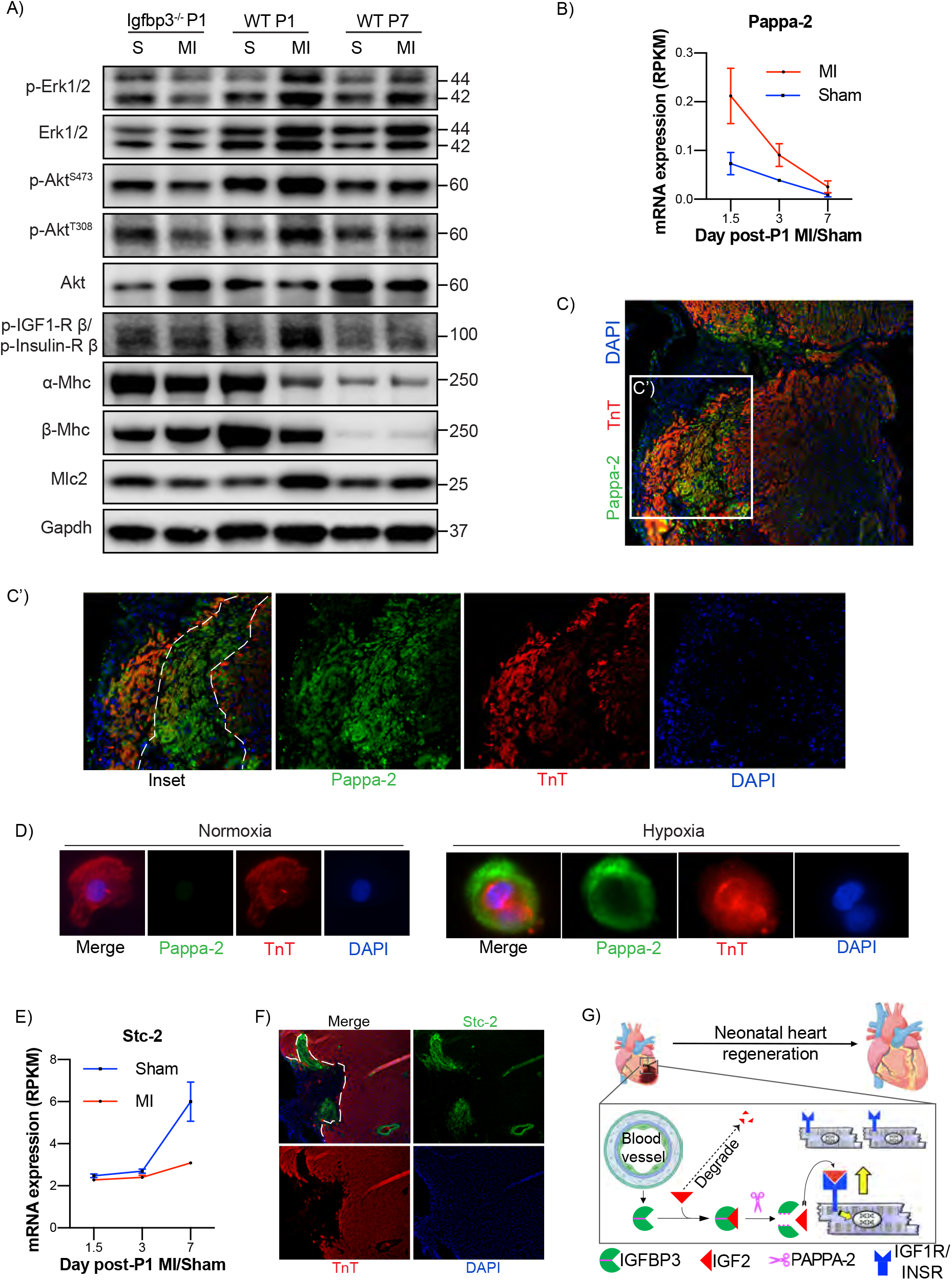
IGFBP3 and its regulators modulate IGF signaling after neonatal MI. A) Western blot of IGF pathway components from cardiac tissue obtained 3 days after regenerating (P1) or non-regenerating (P7) sham or MI injuries. B) Relative mRNA expression (RPKM, reads per kilobase of transcript per million reads mapped) of Pappa2 after P1 MI (red) or sham (blue) operations. C) Epifluorescent microscope images of immunofluorescently-labeled Pappa-2 and TnT cardiac sections 3 days after P1 MI. Dashed white lines demarcate infarct border (C’ is boxed inset of C). D) Epifluorescent microscopic images of NRVM exposed to 21% (normoxia) or 1% oxygen (hypoxia) for 24 hours followed by immunofluorescent antibody staining against Pappa-2 and TnT. E) Epifluorescent microscope images of immunofluorescently-labeled Stc-2and TnT cardiac sections 3 days after P1 MI. Dashed white lines demarcate infarct border. F) Schematic depicting mechanism for IGFBP3-mediated IGF2 stabilization and PAPPA2-mediated release of IGF2 in apposition to cardiomyocytes.

Our findings support a recent study that meticulously characterized IGF2 as the necessary ligand for activation of the cardiomyocyte Insulin Receptor (IR), which effects cardiac regeneration after murine neonatal cardiac injury.^31^ However, this report further uncovered that even though both IR and IGF2 are expressed at the protein level after adult MI, the cardiomyocyte IR is not phosphorylated - indicating that IGF2 within the post-adult injury milieu is not sufficient to activate the IR. This observation hinted at a spectral component that is needed for activation of the cardiomyocyte IR by IGF2. Given our elucidation of the spatial localization of IGFBP3 after neonatal MI - which is not expressed after older MI - we hypothesized that IGFBP3 may be a missing link. We speculated that IGFBP3 could shepherd IGF2 into the anatomic region of tissue loss to generate zones of cardiomyocyte proliferation: coordination and delivery of IGF2 by IGFBP3 may be necessary to activate the IR. However, the binding affinity of IGF2 for IGFBP3 is stronger than for the IR, and thus a mechanism to release IGF2 from IGFBP3 is entailed in this model.^60, 61^

Pappalysin 2 (PAPPA2) is a secreted zinc metalloproteinase with proteolytic activity against IGFBP3 and IGFBP5, which could fragment IGFBP3 and thereby yield free IGF2.^62–65^ Analysis of the neonatal heart RNA-seq demonstrated that Pappa2 expression rises sharply after P1 MI compared to sham 1.5 days after injury, and then slowly decays to sham levels by P8 (Fig. 5B). IF of the heart at P4 after P1 MI showed that PAPPA2 is located predominantly in the infarct zone, with co-expression with cardiomyocyte markers in the adjacent border zone (Fig. 5C). Cleavage of IGFBP3 by PAPPA2 would release IGF2 within the histological site of injury, leading to cardiomyogenesis noted at P8 (after P1 MI) (Fig. 2E-G).^66, 67^ As border zone myocytes express PAPPA2 (based on co-localization), we hypothesized that hypoxia/ischemia may regulate Pappa2 levels. We exposed NRVM to ambient conditions or hypoxia (1% oxygen) and detected intense PAPPA2 immunofluorescent staining in hypoxic NRVM, which also exhibited rounded morphology consistent with previous reports^68^ (Fig. 5E). These results suggest that PAPPA2 is expressed by hypoxic cardiomyocytes, although we cannot exclude coincident expression of PAPPA2 by other cell lineages in the regenerating heart.

Interestingly, PAPPA2 is known to be inhibited by Staniocalcin-2 (STC2), a homodimeric secreted glycoprotein that serves to spatially control PAPPA2 expression and IGF signalling.^69^ We interrogated whether STC2 also contributes to the regulation of IGF2 levels by IGFBP3 and PAPPA2. RNAseq of the neonatal heart showed that Stc2 levels are similar between P1 sham and MI early after injury, but diverge by P8, when Stc2 levels in the P1 MI heart are much lower than in the P1 sham heart (Fig. 5E). Intriguingly, at the tissue level, we also noted localized zones of STC2 expression in the boundaries of the infarct zone and perivascularly at P4 after P1 MI, but not in the infarct zone. (Fig. 5F). These results suggest that, after neonatal MI, overall STC2 expression is significantly curtailed and is spatially limited within the peri-infarct zone, whereas PAPPA2-mediated IGFBP3 cleavage, and IGF2 release, is concomitantly upregulated to elicit cardiomyocyte proliferation via canonical IGF signaling (Fig. 5G).

## Discussion

Neonatal cardiac regeneration is orchestrated by manifold intercommunications among the disparate cell lineages resident in the heart and circulating cells elicited via discrete, heterologous protein-protein interactions. The complexity of this cross-talk has been elegantly enumerated by gene expression profiling of whole cardiac tissue and single cells.^12-15, 70^ Here, we build upon these findings by highlighting a tissue-level pathway of paracrine signaling that illustrates how IGF signaling is regulated in cardiomyocytes along the border and infarct zone to regenerate the neonatal heart.

Our study identified and characterized the role of IGFBP3 as a vascular-derived angiocrine factor that is upregulated after neonatal MI and localized to the area of injury. IGFBP3 is necessary for neonatal regeneration, as its deletion can stymie full regeneration after P1 MI. However, use of a global Igfbp3 knockout model may have led to developmental genetic compensation by upregulation of other IGFBPs, which could have partially rescued the deletion phenotype.^71^ We also expressed rIGFBP3 in mammalian cells and developed an ectopic, cell-specific Igfbp3 transgenic mouse model. In both models, IGFBP3 was sufficient to induce cardiomyocyte mitosis in the older postnatal heart. This is an interesting phenomenon for which we do not have a direct mechanistic explanation, but we postulate that overexpression of IGFBP3 increases the tissue concentration of IGF2 through, followed by release of IGF2 which can be mediated by a number known IGFBP3 proteases.^64, 65, 72-76^ In support of our observation, a previously generated global Igfbp3 overexpression transgenic mouse model was also noted to have multi-organ enlargement, including the heart, although this phenotype was not further scrutinized.^77^

As a paracrine cardiomyocyte pro-mitotic factor, IGFBP3 belongs to a class of soluble proteins that have long been studied for their potential to renew the adult mammalian heart. Tissue-resident non-myocyte lineages that elaborate secreted proteins with cardiomyogenic capacity include epicardial cells (FGFs, FSTL1, IGFs), endothelium (FGFs, NRG1), and fibroblasts (fibronectin, collagen, HBEGF).^31, 78-82^ Circulating cells have also been found to secrete candidate mitogens in the heart, including macrophages (CCL24, CLCF1) and T cells (CCL24, AREG, GAS6).^13, 14, 83^ These proteins have been discovered from studies of cardiac development as well as from postnatal contexts (including neonatal regeneration.) In addition, the mechanism of action of some of these factors has recently been defined. For example, NRG1 binds to the heterodimeric ERBB2/ERBB4 tyrosine kinase receptor on the cardiomyocyte plasma membrane to promote cell-autonomous dedifferentiation and proliferation. However, the ability of NRG1 to stimulate myocyte proliferation is limited to the neonatal period, as older cardiomyocytes cease to express ERBB2; ERBB4 homodimers cannot initiate downstream mitogenic pathways.^84–86^

The prominent activity of the IGF ligands during cardiac development in zebrafish and mice is well-established.^32, 67, 87, 88^ Moreover, characterization of embryonic stem cell-derived cardiomyocytes and murine neonatal regeneration also evinced the principal role of the IGF signaling pathway in cardiomyocyte proliferation during in vitro and postnatal environments.^31, 89^ However, the purported salutary effect of IGF ligands on cardiomyocytes in the adult heart did not materialize in animal studies.^34, 35, 90^ Hence, IGF pathway modulation-based mechanisms have largely been eschewed over the last decade. Our identification of an IGFBP3-PAPPA2-STC2 axis that regulates and coordinates IGF2 activity post-translationally may reconcile some of these previous disparate studies. Indeed, IGFBP3 is not itself a mitogen but it helps conduct neonatal heart regeneration within the auspices of the canonical IGF pathway. IGFBP3 may concentrate IGF2 levels in a site-specific manner within the injury zone to activate its cognate receptor, which cannot occur in the adult heart due to the absence of IGFBP3 and the diffusibility of the 7.4 kDa mature IGF2 protein.^30, 31^ Although we mapped the spatial and temporal expression of PAPPA2 and STC2 in the regenerating heart, which highlights the importance of spatial coordination among these various effectors to modulate local IGF signaling at the injury zone, we cannot exclude the possibility that other IGFBP3 proteases may also play a role in the release and delivery of IGF2.^35, 72-74^

While the IGF binding proteins can exhibit Janus-like effects through IGF-dependent (extracellular) and IGF-independent (intracellular) activities in different cell systems and disease states, we elucidate here that IGFBP3 acts as an upstream regulator of IGF pathway during neonatal cardiac regeneration. Our study provides insights that add to the growing list of pro-regenerative pathways that differentiate the neonatal and adult hearts. Our data suggests that STC2, by restraining PAPPA2 activity and promoting IGF2 sequestration by IGFBP3, may play a role in the decline in cardiomyocyte proliferation with age – which merits further exploration. Overall, our model highlights the complexity of the spatial regulation of cardiomyocyte proliferation and supports the potential role of IGFBP3 as a pro-regenerative cardiomyogenic factor.

## Acknowledgements

We thank Dr. Vincent Tagliabracci and lab members for helpful discussions and providing reagents. We are grateful to Dr. Hao Zhu and Dr. Tripti Sharma and the UTSW CRI Transgenic Core for assistance with generating the Igbp3 overexpression transgenic mouse model. We thank Dr. Robert Baxter for helpful discussions. S.R.A. was supported by NIH 5T32HL125247-03 and NIH K08HL153788. H.A.S. was supported by NIH R01 HL137415-02, NIH R01 HL147276-01, NIH R01 HL149137-01 and Leducq Transatlantic Network of Excellence

## Methods

### Mice

All mouse experiments were performed as per protocols approved by the Institutional Animal Care and Use Committee (IACUC) at the University of Texas Southwestern Medical Center (UTSW). All animal experiments complied with relevant ethical regulations on animal research. Mice had *ad libitum* access to water and food and were housed in 12:12 hour light:dark cycles in a temperature-controlled room in the Animal Research Center at UTSW. The age of the animal is indicated in the text and/or in the figure legend for each experiment. Littermate controls were used whenever possible for experiments with multiple genotypes. Statistical tests were not used to predetermine sample size. All surgeries and echocardiographic studies were carried out blinded to the genotype of the mice during the experiments and outcome assessments. CD1 mice (from Charles River Laboratories) were used as wild-type mice for studies. The Igfbp3^-/-^ and Igfbp3 LacZ mice are equivalent and obtained from the Knockout Mouse Project.^36^ The Igfbp3^OE^ mouse model was generated with assistance from the UTSW Cancer Research Institute (CRI) Transgenic Core Facility (description below). Tie2-CreERT2 and Cdh5-CreERT2 mice have been previously described.^58, 59^

### Drug Administration

Tamoxifen (Sigma) was prepared by dissolution in sesame oil (Sigma) to a concentration of 20 mg/mL. 4-hydroxytamoxifen (4-OHT) (Sigma) was prepared by dissolution in ethanol (10%) and sesame oil (90%) at a concentration of 1 mg/mL. Both TM and 4-OHT were administered by intraperitoneal injections. Cre induction for Tie2^OE^ mice in the neonatal period was accomplished with 125μg 4-OHT on P1, 250μg 4-OHT on P3, and 1mg TM on P5; the mice were euthanized on P7 (Fig. 4). Cre was induced for the Cdh5^OE^ mice (Fig. 4) with 0.5 mg TM at P28 and P30, 1mg TM on P32 and P33, and 2mg TM on P37-P39 (daily). The mice were euthanized 2.5 weeks after the last injection. Mice in an experiment were euthanized at the same approximate time of day to avoid circadian variability.

### Mouse model of neonatal MI

Neonatal anterior wall MI was performed as previously described.^91^ In brief, 1-day-old pups were anesthetized by hypothermia on ice for 3 to 5 minutes, then placed on a cooled platform in a right lateral position to expose the left thorax. Following lateral thoracotomy, the LAD coronary artery was identified along the LV anterior wall when possible. Prolene sutures (6-0) were used to tie off the LAD to induce infarction. If the LAD was not visible, then ligation was performed on the expected anatomic location. Blanching of the myocardium below the ligature was observed as an indicator of ischemia. Then, 6-0 nonabsorbable Prolene sutures were used to suture the ribs together and seal the chest wall incisions. Skin glue was used to join the skin together. The pups were then warmed rapidly under a heat lamp till arousal and urine output was achieved. The pups after surgery were placed on a heat pad until the entire litter had undergone surgery, at which point the mice were mixed with bedding from the mother’s cage and placed back with the mother (or foster, if the litter was the mother’s first). The pups were euthanized at noted time points for analysis, including 28-30 days after MI (or sham) for cardiac function analysis with echocardiography. Fibrotic scar size was quantified using ImageJ on five Picorisius Red-stained sections one month after P1 MI below the level of ligature per heart.

### Transthoracic echocardiography

Assessment of cardiac function was performed on conscious, non-sedated mice using a Vevo2100 micro-ultrasound system, MS400C probe (VisualSonics) one month after P1 MI. Echocardiographic M-mode images were obtained from a parasternal short-axis perspective at the level of the papillary muscles. LV internal diameters at end-diastole (LVIDd) and end systole (LVIDs) were determined by M-mode images. Six representative contraction cycles were selected for analysis, and average indices (LVIDs, LVIDs, EF, and FS) were determined for each mouse. All echocardiography measurements were performed by a blinded operator.

### Histology

The hearts were collected and treated in 4% paraformaldehyde fixative (in PBS) overnight at 4°C and then processed for either paraffin or cryo embedding. H&E and Picosirius Red staining were performed according to standard procedures at UTSW Histology core facility on paraffin sections.

### LacZ Staining

Whole mount LacZ staining was performed after fixing the freshly-isolated hearts in 0.2% glutaraldehyde for 3 hours at 4°C. The hearts were then washed in detergent rinse solution (0.02% Igepal, 0.01% Sodium Deoxycholate, and 2mM MgCl_2_ in 0.1M phosphate buffered saline (pH 7.4) three times, then incubated in X-gal staining solution (0.02% Igepal, 0.01% Sodium Deoxycholate, 5mM Potassium Ferricyanide, 5mM Potassium Ferrocyanide, 2mM MgCl_2_, 1mg/ml X-Gal (from stock prepared in N,N-dimethylformamide) diluted in 0.1M phosphate buffered saline (pH 7.4)) for 48h at 4°C. The hearts were then washed in PBS and post-fixed in 4% PFA overnight at 4°C, then imaged and stored or processed for paraffin sectioning and Nuclear Fast Red counterstain.

### Immunofluorescence staining

Immunostaining was performed according to prior reports. Briefly, heart cryosections were equilibrated with antigen retrieval buffer in epitope retrieval buffer (IHC World) or 1x citrate buffer (Antigen Retrieval Citra Plus, Biogenex). Samples were permeabilized and blocked with 0.3% Triton X-100 and 10% serum from the host animal of secondary antibodies in PBS for 1 hour at room temperature. Then, the samples were incubated overnight at 4°C with primary antibodies. After three washes in PBS, samples were incubated at room temperature for 1h with the corresponding fluorescence secondary antibodies conjugated to Alexa Fluor 488 or 555 (Invitrogen) at 1:400. The slides were mounted in Vectashield Antifade Mounting Medium (Vector Laboratories). Slides were viewed under Nikon fluorescence or Zeiss LSM 510 confocal microscopes. Primary antibodies used were: phosphohistone H3 Ser10 (EMD Millipore, 06-570; 1:100), aurora B kinase (Sigma, A5102; 1:100); troponin T, cardiac isoform Ab-1, clone 13-11 (Thermo Scientific, MS-295-P1; 1:200), sarcomeric α-actinin (Abcam, ab68167; 1:200); flag (Genescript, A00187-100; 1:200); endomucin (Santa Cruz, V.7C7; 1:1250); Pappa2 (Thermo Scientific, PA5-21046; 1:100); Stc2 (Abcam, ab63057; 1:100); Igfbp3 (Santa Cruz, sc-9028; 1:100). DAPI or Hoechst was used for nuclear staining. Images were obtained on a Nikon Eclipse Ni or Nikon A1 laser scanning confocal microscopes.

### WGA staining and cardiomyocyte size quantification

WGA staining and quantification was performed as per prior report.^92^ Briefly, the slides were incubated with WGA conjugated to Alexa Fluor 488 (50 mg/mL, Life Technologies) for 1h at room temperature following PBS washes. To quantify the cross-sectional cardiomyocyte cell size, three to four independent hearts per genotype/group were captured at 40x magnification from three different views and positions (e.g. right ventricle, left ventricle, septum). Cellprofiler was used to quantify the size of cardiomyocytes were round and had a nucleus.^93^ Quantification of at least 500 cells per sample was performed.

### Cardiomyocyte isolation

Adult hearts were freshly collected and fixed in 4% PFA at 4°C overnight after removal of atria. The heart was cut from the apex towards the base twice (in perpendicular cuts) while preserving the basal connections to expose the inside surfaces of the heart. The hearts were subsequently incubated with collagenase type 2 (Worthington-Biochem, Cat# LS004176) supplemented with 1% penicillin-streptomycin (Thermo Fisher) overnight at 37°C with constant rotation. For the next 7 days, the supernatant was collected twice a day and stored at 4°C (supplemented with fetal bovine serum (Hyclone)) while additional collagenase 2 was added to the remnant cardiac tissue for further digestion. At the end of the 7-day period, the cardiac tissue was mostly connective tissue, and the pooled cells were allowed to settle by gravity before aspiration of the supernatant followed by resuspension in 1 mL of PBS and passage through a 150-μm nylon mesh filter. The isolated cardiomyocytes were then counted after serial dilution using a bright field microscope.

### Western blotting

Ventricles were collected and lysed in RIPA buffer with the addition of complete protease inhibitor cocktail (Roche). Protein concentration was determined using Pierce BCA protein assay kit (Pierce Biotechnology), with three biological replicates. Following separation via SDS-PAGE gels, proteins were transferred to nitrocellulose membranes (Bio-Rad), blocked in 5% skim milk/TBST (TBS with 0.1% Tween-20), and incubated with primary antibodies: Igfbp3 (Santacruz), Flag (Genscript), anti-p44/42 MAPK (Erk1/2) (Cell Signaling, 9102S, 1:500), anti-phospho p44/42 MAPK (Erk1/2) (Thr202/Tyr204) (Cell Signaling, 9101S, 1:2000), anti-Akt (Cell Signaling, 9272S, 1:1000), anti-phospho Akt (Thr308) (Cell Signaling, 9275S, 1:1000), anti-phospho Akt (S473) (Cell Signaling, 4060S, 1:1000), anti-Myosin Light Chain 2/MLC-2V (Proteintech, 10906-1-AP, 1:1000), anti-MyHC-α (Sigma, HPA001349, 1:3000), anti-MyHC-β (Sigma, HPA001239, 1:2000), anti-phospho-IGF-I Receptor β (Tyr1135/1136)/Insulin Receptor β (Tyr1150/1151 (Cell Signaling, 3024S, 1:1000), anti-Gapdh (Sigma, AB2302, 1:5000). Horseradish peroxidase-conjugated peroxidase anti-mouse, antirabbit, or anti-goat antibodies (Trublot, Rockland (18-8817-30, 18-8816-31, 18-8814-31; 1:5000) or Jackson ImmunoResearch (115-035-166, 111-035-144, 703-035-155, 705-035-147; 1:25,000–1:50,000) were used as secondary antibodies. The membranes were exposed using Licor Odyssey Fc system and quantified by Image Studio Lite v.5.2 software.

### IGFBP3-Flag plasmid cloning, protein expression and purification

Mouse IGFBP3 cDNA fused with a C-terminal 3xFlag (linked with a glycine-glycine-glycine-glycine-serine segment) was cloned into pQXCIP (Clontech) retroviral vector and sequence-confirmed. 293T cells were co-transfected with 5μg of the Igfbp3-Flag plasmid and 5μg of pCL-10A1 (Novus Biologicals) using Lipofectamine 3000 (Invitrogen). 48h later, polybrene at final concentration of 8μg/mL was added. 48h later, selection with puromycin at final concentration of 1μg/mL was begun for 14 days. Six independent single-cell clones were later selected and validated by qPCR (Igfbp3) and Western blot (Flag) as well as IGFBP3 levels in the conditioned medium (using ELISA). One clone with the highest levels of secreted IGFBP3-flag was selected for further experiments. To purify protein, conditioned medium from five 10-cm plates was pooled, spun at 300g to remove cells, and incubated with 200 μL of M2 Flag Agarose beads (Sigma) overnight at 4°C. The next day, the beads were magnetically separated and washed in wash buffer (50 mM Tris, 150 mM NaCl, pH 8.0) three times, then resuspended in 100 μL of buffer (50 mM Tris, 150 mM NaCl, 1mM EDTA, pH 7.4) with Flag peptide (Sigma) and rocked overnight at 4°C overnight. The next day, the beads were magnetically pelleted and the supernatant was collected and confirmed by Western blot (Flag) and concentration assessed by BCA assay (Pierce). All steps included protease and phosphatase inhibitors (Thermo Scientific). PNGase F (NEB) treatment was performed on 1μg purified and denatured IGFBP3-flag protein according to the manufacturer’s instructions.

### Cre lentivirus production

293T cells were plated until 70% confluence on 10-cm plates, at which time they were transfected with 5μg of Cre-ire-puro vector (Addgene #30205) along with 3μg pAX2 and 2μg VSVG packaging plasmids using Lipofectamine 3000 (Invitrogen). 48 hours later, the supernatant was collected and spun at 300*g* for 5 minutes to remove cells and debris. The resulting lentivirus-laden supernatant was filtered through a 0.45 μm polyethersulfone membrane filter (Thermo Scientific 725-2545) and stored at −80°C until use.

### Generation of primary ear tip fibroblasts

2-3 ear slices (~25 x 25 mm) were obtained from mice with clean scissors and rinsed in 100% ethanol followed by PBS. Slices for a single mouse were then coated in 100% Matrigel and placed in one well of a 6-well plate, followed by incubation at 37°C for 30 minutes. Then, medium was gently added to the well to cover the ear slide/Matrigel mixture and incubated at 37°C; medium consisted of DMEM-F12 (Thermo Fisher Scientific) with 10% FBS and 20 ng/mL FGF (Merck Millipore). Fresh medium containing FGF was added every 2 days. After 80-100% confluence was reached, fibroblasts were passaged to a 6-well plate for further experiments. For Cre lentivirus infection, cells were grown to 60% confluence in a single well of a 6-well plate, then the medium was removed and replaced with 1mL of Optimem (Thermo Fisher Scientific) supplemented with 16 μg/mL polybrene as well as 1 mL of the Cre lentivirus supernatant. 24h later, the medium was replaced with regular maintenance medium. 48h after lentiviral transduction, 1 μg/mL puromycin-containing medium was added for up to two weeks to select for infected cells. Lentivirus without Cre was used control. For ELISA measurements of conditioned medium for tail tip fibroblasts, the conditioned medium after 2 weeks in puromycin selection was collected and spun at 300*g* to remove cells and debris, followed by performance of IGFBP3 ELISA (R&D Systems) on the supernatant according to manufacturer’s instructions.

### Generation of Igfbp3-overexpression conditional mouse model

Mouse Igfbp3 cDNA (Origene) was amplified with attB-containing primers and cloned into pDONR 221 (Invitrogen) using BP Clonase (Invitrogen) in One Shot Top10 chemically competent cells (Invitrogen). Proper insertion was confirmed by sequencing. Then, the resulting plasmid and pR26 CAG/GFP Dest plasmid (Addgene# 74821) were Gateway-recombined using LR Clonase (Invitrogen) to generate the Rosa26-CAG-Stop-LoxP-Stop-Igfbp3-ires-GFP plasmid. The plasmid was sequence-confirmed and linearized prior to co-microinjection into zygotes with Cas9-bearing plasmid as previously described.^54^

Specifically, Alt-R S.p. Cas9 Nuclease V3 (Integrated DNA Technologies), sgRosa 26 - ACUCCAGUCUUUCUAGAAGA (Integrated DNA Technologies), TracrRNA (Integrated DNA Technologies) and the recombineering plasmid Rosa26-CAG-Stop-LoxP-Stop-Igfbp3-ires-GFP plasmid were microinjected into C57BL/6J zygoes. Injected zygotes were transferred into the oviducts of pseudo-pregnant CD1 female mice to obtain live pups. Chimeric mice were genotyped by polymerase chain reaction (PCR) analysis to confirm knock-in into the Rosa26 locus.^54^ Founder mice were backcrossed with C57BL/6J mice for three generations prior to crossing with Cre-bearing mice. Male Cre mice were crossed with female Igfbp3-overexpression mice to obtain experimental animals.

### Cardiomyocyte In Vitro Proliferation Assessment

NRVMs were derived from 1- or 2-day old Sprague-Dawley rates using the Cellutron Neomyt kit (Thermo Fisher Scientific). Cell purity was routinely >90% cardiomyocytes. These cells were plated at a density of ≈800 cells/mm^2^ to enrich for cardiac myocytes and cultured for 24 hours in DMEM/M199 (3:1 ratio) containing 5% fetal bovine serum, 10% horse serum, and 100 μM bromodeoxyuridine (Sigma-Aldrich) on poly-lysine-coated coverslips at 37°C. Subsequently, the medium was replaced with conditioned medium from the Igfbp3-flag-expressing 293T clone (described above) 2 days after removal of puromycin-containing medium. 48h after incubation in the respective medium, the coverslips were washed in PBS three times, then fixed in 4% PFA for 15 minutes at room temperature, then blocked in 10% goat serum with 0.1% Triton X-100 for 1h at room temperature, then incubated in primary antibodies (Cardiac Troponin T (Thermo Scientific) and pH3 (EMD Millipore) prepared at 1:100 dilution in 3% goat serum with 0.3% Tween-20) overnight at 4°C. The following day, the coverslips were washed and incubated at room temperature in fluorescent species-specific Alexa Fluor-conjugated antibodies (Invitrogen) at 1:400 for 1h. This was followed by series of washes and treatment with DAPI (Invitrogen), followed by mounting in Vectashield. Three replicates for each group were imaged in 6 high-powered field images per sample on a Nikon microscope. The total number of cardiomyocytes (based on cardiac Troponin T expression) and the total number of pH3+ cardiomyocytes (based on cardiac Troponin T and nuclear pH3 co-expression) were assessed.

### Cell culture

Cultured cells were maintained at 37°C in a humidified incubator with 5% CO2. 293T cells were cultured in Dulbecco’s modified Eagle’s medium (DMEM;Invitrogen) containing 10% (v/v) fetal bovine serum (FBS; Hyclone) and 1% (v/v) penicillin/streptomycin (Thermo Fisher Scientific).

### Co-immunoprecipitation

NRVM were incubated with IGFBP3-flag-containing conditioned medium (or wildtype conditioned medium for control) from 293T cells for 24h, as described above. Cells were washed in cold PBS and protein harvested in IP lysis buffer: 2.5mM Tris, pH7.4, 150 mM NaCl, 1 mM EDTA, 1% NP40, 5% glycerol. Protease inhibitor cocktail and phosphatase inhibitor were included at all times. Protein concentrations were determined by BCA assay (Pierce) per manufacturer’s instructions. 500 μg of protein from each group was incubated with 50 μL of M2 Flag Agarose beads (Sigma) overnight with gentle rocking at 4°C. After magnetic separation, the pellets were washed three times with 500 μL IP lysis buffer. The pellets were resuspended in 250 μL of IP lysis buffer with addition of Flag peptide (Sigma), and the mixture was rocked overnight at 4°C. The following day, magnetic separation was used to collect the supernatant. The supernatant was mixed with 50 μL 4XSDS sample buffer and boiled for 10 minutes. Then, 50 μL of the sample was loaded onto SDS-PAGE gel (4-20%). When samples entered the resolving gel for about 5-10 mm, the run was stopped and the gel was stained with Coomassie Blue. The stained area was cut and diced into 1mm cubes. The samples are then subjected to mass spectrometer analysis for protein identification.

### Statistics and reproducibility

Differences between groups were examined using unpaired two-sided Student’s t-test to determine statistical significance. All bar graphs represent mean +/- s.e.m. *p* values are shown in graphs. The sample sizes were as follows: Fig. 1B-C, n=3 per group; Fig. 1D, n=3 per group; Fig. 1F-J, n=3 per group; Fig. 1K-N, n=3; Fig. 1O, n=3; Fig. 1P, n=3); Fig. 2B, n=3 per group; Fig. 2C, n=3 WT, n=14 Igfbp3^+/-^, n=12 Igfbp3^-/-^; Fig. 2D, n=3 WT, n=10 Igfbp3^+/-^, n=7 Igfbp3^-/-^; Fig. 2E, n=3; Fig. 2F, n=6 per group; Fig. 2G, n=3 per group; Fig. 2H, n=3 per group; Fig. I-K, n=10 WT, n=15 Igfbp3^+/-^, n=12 Igfbp3^-/-^, Fig. 2L-M, n=3 WT, n=6 Igfbp3^-/-^; Fig 3B, n=3 per group; Fig. 3C, n=3 per group; Fig. 3D, n=2; Fig. 3E, n=2; Fig. 3F-G, n=8-9; Fig. 3H, n=3; Fig. I-J, n=3; Fig. 4B, n=3; Fig. 4C, n=3; Fig. 4D-E, n=3 control, n=5 OE; Fig. 4F, n=3; Fig. 4G, n=3 control, n=3 OE; Fig. 5A, n=3; Fig. 5B, n=3; Fig. 5C-D, n=3; Fig. 5E, n=3. Fig. S1, n=3; Fig. S2A, n=4 WT, n=3 Igfbp3^-/-^; Fig. 5B, n=3 WT, n= 7 Igfbp3^+/-^, n=9 Igfbp3^-/-^; Fig. S3A-B, n=3; Fig. S4A, n=2; Fig. S4D, n=3; Fig. S4E, n=2; Fig. S5, n=3.

### Data availability

The data that support the findings of this study are available within the paper and its Supplementary Materials. Source data or other materials are available from the corresponding authors upon reasonable request.

---

**Supplementary Figure S1**: Validation of microarray screen results. A) In-situ hybridization for Angiopoietin-like 7 (Angptl7) and TnT 3 days after P1 MI (top panel) or P7 MI (bottom panel), with associated insets.

**Supplementary Figure S2**: Characterization of Igfbp3^-/-^ mice. A) Systolic function of uninjured 1-month-old mice (EF). B) HW/BW ratios (mg/g) for 1-month-old mice after P1 MI.

**Supplementary Figure S3**: Validation of IGFBP3 antibody in tissue. A) Immunofluorescent labeling of IGFBP3 antibody on Igfbp3^-/-^ tissue 3 days after P1 MI using epifluorescence microscopy (scale bar 100 μm). B) Secondary antibody only staining of WT cardiac tissue 3 days after P1 MI using epifluorescence microscopy (scale bar 100 μm).

**Supplementary Figure S4**: In vitro and in vivo models of ectopic IGFBP3. A) Western blot for FLAG from conditioned medium of IGFBP3-overexpression 293T cells (lanes 1-5) or WT cells (lane 6) with different treatment durations of tunicamycin. B) Genomic DNA PCR for knock-in construct of CAG-Igfbp3 cassette demonstrating recombination into the native Rosa26 locus. C) Immunostaining for TdTomato (TdT) in the Tie2CreERT2;R26^loxp-stop-loxp-TdT^ (Ai14) mouse model.

**Supplementary Figure S5**: Western blot quantification from Figure 5. All values obtained on Western blot derived from ventricular tissue obtained 3 days after the annotated operation. A) Relative densitometry of pERK1/2 to ERK1/2. B) Relative densitometry of pERK1/2 to GAPDH. C) Relative densitometry of pAKT-S473 to total AKT. D) Relative densitometry of pAKT-S473 to GAPDH. E) Relative densitometry of pAKT-T308 to total-AKT. F) Relative densitometry of pAKT-T308 to GAPDH. G) Relative densitometry of pIGF1R/pINSR to GAPDH. (S= sham).

**Supplementary Table 1**: Microarray screen results for predicted secreted genes (Figure 1a).

**Supplementary Table 2**: Proteomics results after coimmunoprecipitation of Flag from protein lysate derived from NRVM with IGFBP3-flag-conditioned medium (or wildtype conditioned medium).

