## Supplemental Figures for "Angiocrine IGFBP3 Spatially Coordinates IGF Signaling During Neonatal Cardiac Regeneration"

Supplementary Figure S1: Validation of microarray screen results

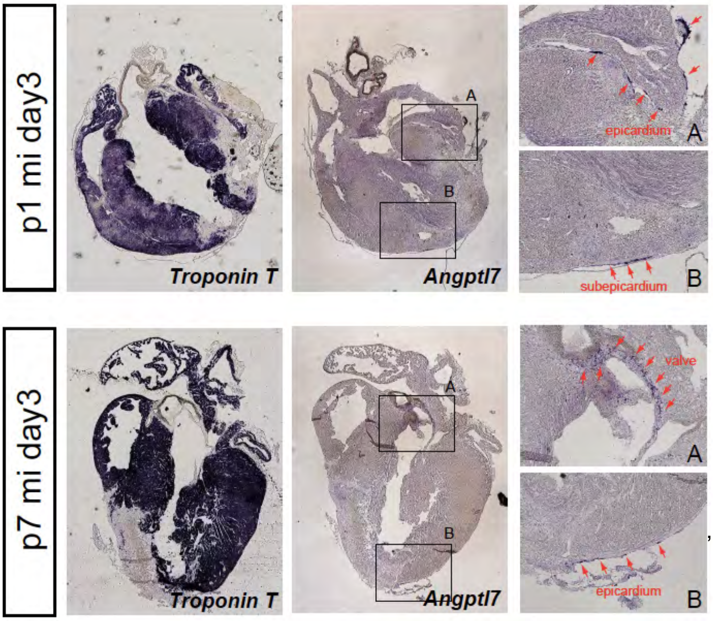


,

,

,

,

Supplementary Figure S2: Characterization of Igfbp3-/- mice.

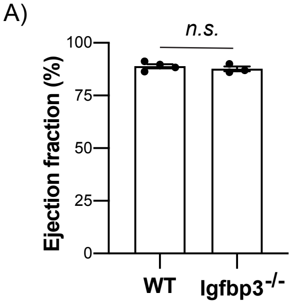


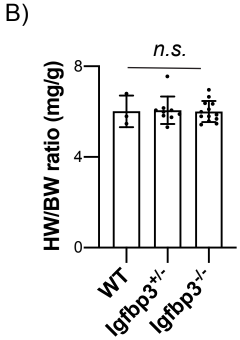

Supplementary Figure S3: Validation of IGFBP3 antibody in tissue.

A)

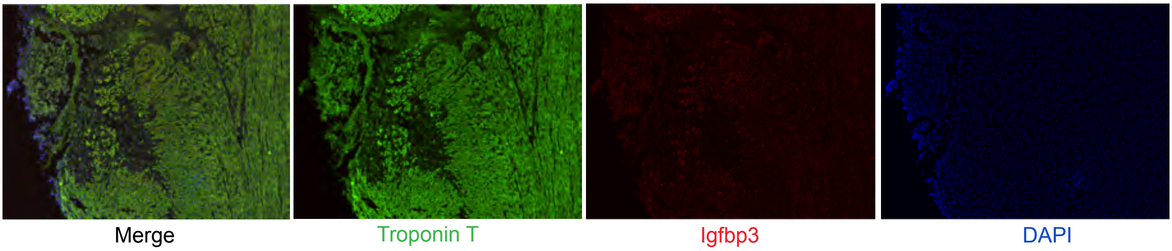


B)

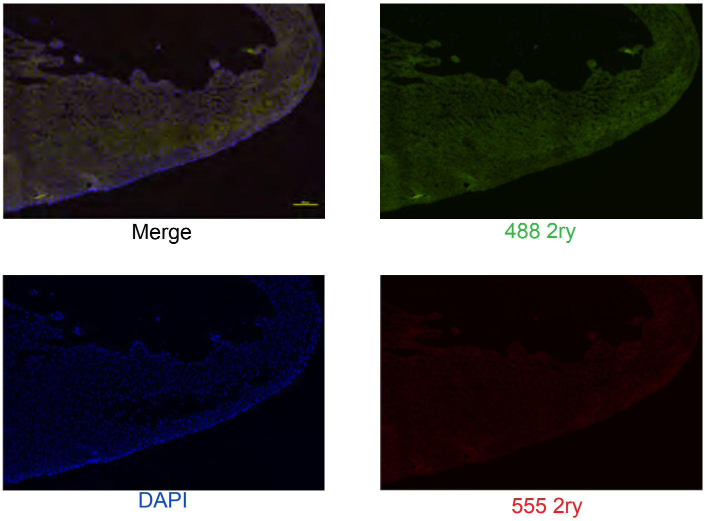

Supplementary Figure S4: In vitro and in vivo models of ectopic IGFBP3

A)

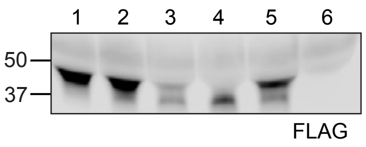


| | Cell type | Tunicamycin | Media change upon tunicamycin |
|---|---|---|---|
| Lane 1 | Igfbp3-flag-OE | No | N/a |
| Lane 2 | Igfbp3-flag-OE | 8h treatment | No |
| Lane 3 | Igfbp3-flag-OE | 8h treatment | Yes |
| Lane 4 | Igfbp3-flag-OE | 24h treatment | Yes |
| Lane 5 | Igfbp3-flag-OE | 24h treatment | Yes |
| Lane 6 | WT | No | N/a |

B)

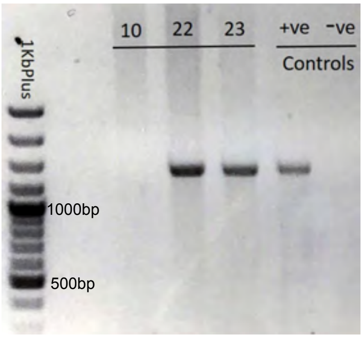


C)

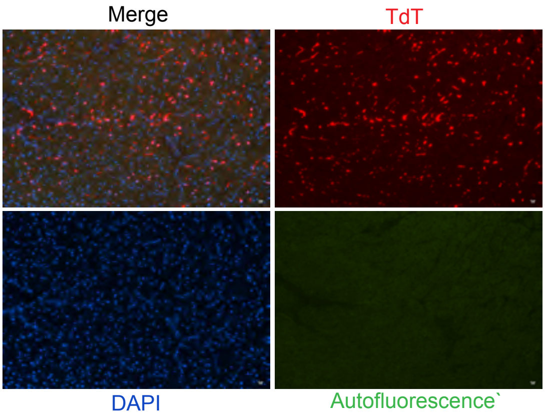

Supplementary Figure S5: Western blot quantification (Figure 5)

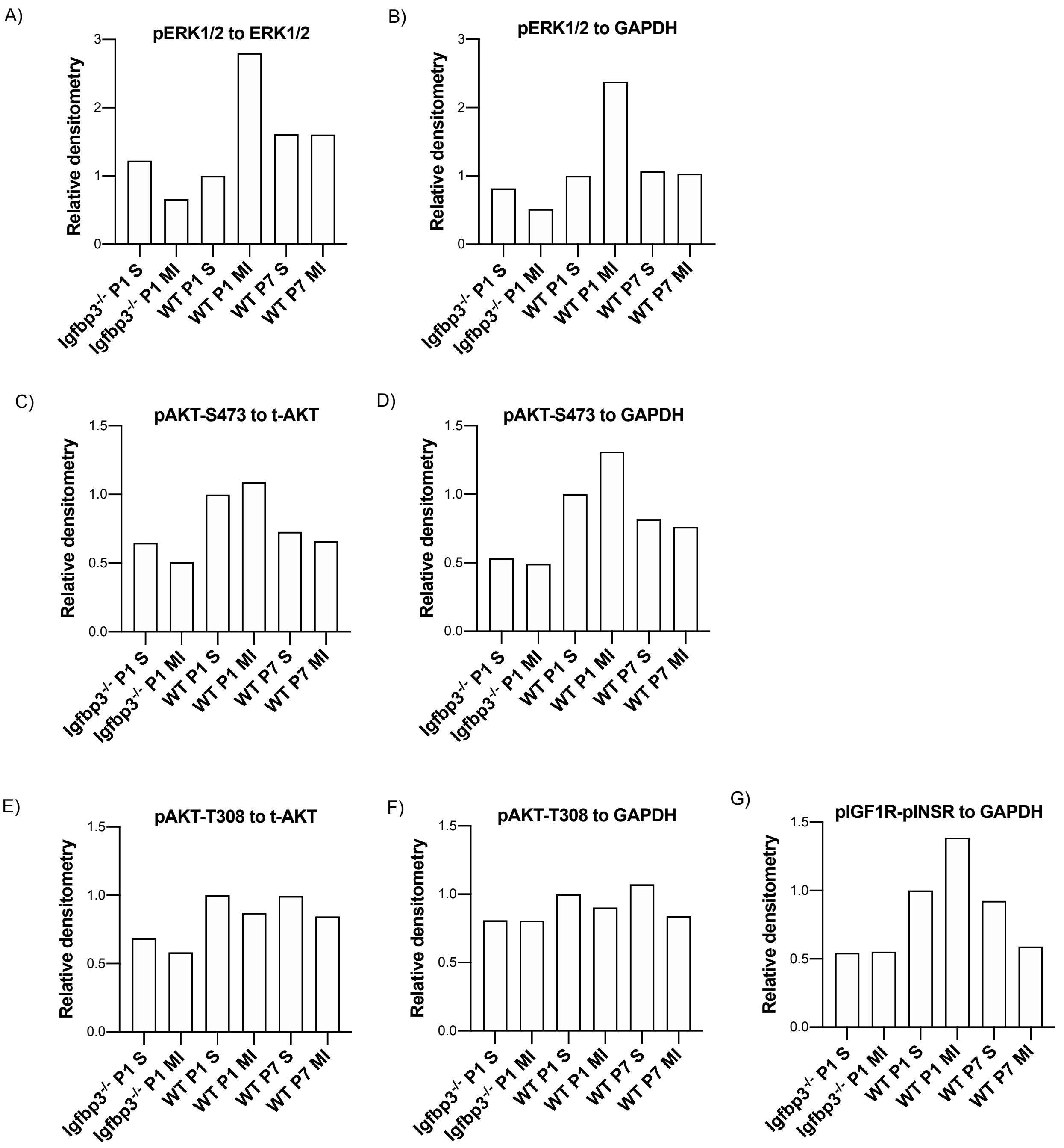
